# A role for CD4 helper cells in HIV control and progression

**DOI:** 10.1101/2021.12.20.473599

**Authors:** Igor M. Rouzine

**Affiliations:** Laboratory of Computational and Quantitative Biology, Institut Biologie Paris-Seine, Sorbonne Université, Campus Pierre et Marie Curie, Paris 75005; Immunogenetics Group, Sechenov Institute of Evolutionary Physiology and Biochemistry, Russian Academy of Sciences, Saint-Petersburg 194223

## Abstract

It remains unclear why HIV persists in most untreated individuals, and why a small minority of individuals can control the virus, either spontaneously or after an early treatment. The present work motivated by the striking differences in the functional avidity of CD4 T cells discovered between patient cohorts in a recent study [1] offers an experimentally–testable mathematical model that explains the diverse outcome of infection. The model predicts an arms race between viral dissemination and the proliferation of HIV-specific CD4 helper cells leading to one of two states: a low-viremia state or a high-viremia state. Helper CD4 cells with a higher avidity favor virus control. The parameter segregating spontaneous and post-treatment controllers is the infectivity asymmetry between activated and resting CD4 T cells. The predictions are found to be consistent with the data from [1] and with data on the avidity CD8 T cells [2]. I also analyze the alternative explanation of T cell exhaustion previously proposed to explain the diverse patient cohorts and demonstrate that it does not explain these and some other experimental data.

**Importance:** Why HIV persists in most untreated individuals, and why a minority can control the virus, either spontaneously or after short treatment, remains unknown. The present study offers a mathematical model of the immune response that explains the differences between the patient cohorts as a result of the arms race between viral dissemination and the proliferation of HIV-specific CD4 T cells. It offers testable predictions and personalized adjustment of early ART to a patient.

## Introduction

Untreated HIV infection causes a gradual loss of CD4 T cells that eventually causes AIDS. Among possible reasons of the slow depletion are impaired homeostasis [3] and a viral adaptation to the host [4]. However, a small fraction of individuals is capable of controlling HIV replication even in the absence of antiretroviral treatment (ART) [5, 6]. These “controllers” exhibit low viremia undetectable by standard assays, far below their levels in patients receiving ART [7-9]. Importantly, HIV controllers very slowly progress to AIDS if at all [7], confirming the link between a low viral load and the delay of AIDS symptoms [4, 10].

Substantial evidence demonstrates that controllers suppress HIV replication due to an active immune response [11]. CD8 T cells rapidly kill infected CD4 T cells through cytotoxic mechanisms involving perforin and Granzyme B [12, 13]. The signs and markers of immune activation are much more pronounced in HIV controllers than in ART patients, including immunoproliferation rate [14], markers of T cell activation [15], and the increased secretion of cytokines IFN-γ and MIP-1β after stimulation with polyclonal antibodies [16]. While ART patients with undetectable viral loads show a gradual fading of the cellular immune responses [17], HIV controllers maintain polyfunctional effector memory T cells that can secrete multiple cytokines [18-20].

Genetic studies show that the variation of HLA subtypes of CD8 T cell receptors among individuals remains the strongest genetic determinant of HIV progression [21-24]. An experimental study demonstrated that HLA-B57, the allele the most associated with HIV control, has unusually broad cross-recognition of antigenic variants of virus [25]. A number of studies compared between patient cohorts the functional avidity, a parameter quantitating the ability of T cells to respond to antigen. In some studies, CD8 T-cell responses in HIV controllers were associated with a high avidity of CD8 T cells [26-29], whereas the other studies indicated a similar avidity in progressors and controllers [30-34]. A recent work concluded this debate demonstrating that that the range of the functional avidity of HIV-specific CD8 T cells does not differ much between a progressive and non-progressive infection [2].

A striking feature setting the two cohorts apart is an active CD4 T cell response with IL-2 secretion capacity and proliferative responses observed in controllers but not in progressors [16]. In a recent study, the functional avidity of Gag293-specific CD4 T cells was compared between spontaneous controllers, high viremia patients, and patients who underwent highly active ART [1]. The avidity was evaluated by IFN-γ ELISPOT assay in the presence of decreasing peptide concentrations with serial dilutions from 4 × 10^−6^ M to 10^−11^ M. For each peptide dilution, IFN-γ production was expressed as the number of spot-forming cells per million cells. The functional avidity was defined as the last peptide dilution that gave a positive IFN-γ response 2-fold above the background level or higher. HIV controllers contained memory CD4+ T cells that differentiated upon antigen stimulation into effector CD4+ T cells with a generally higher functional avidity for an immunodominant Gag epitope than in progressors, consistent with a higher binding affinity of TCR to peptide/MHC complex [1]. Although the avidity in all cohorts of patients was broadly distributed, the distribution for controllers was narrower than for progressors and centered at much higher values (Fig. 1). The explanation for these experimental results is absent.

**Fig. 1.**
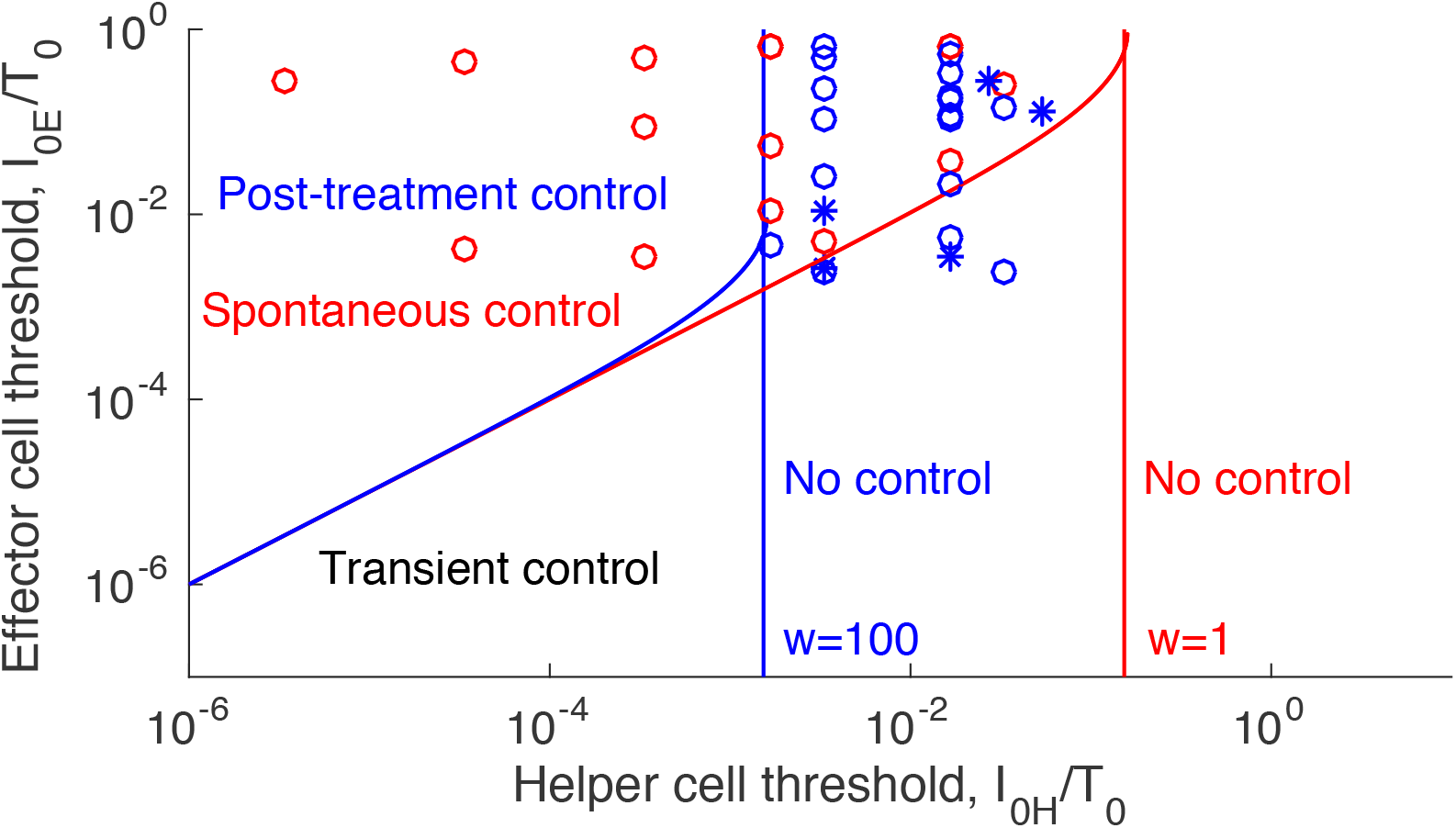
The difference in CD4 cell avidity between controllers and progressors based on. [1]. X-axis: Rescaled inverse functional avidity of CD4 T cells, denoted *I*_*H*0_, measured in three different cohorts of patients [1]. Y-axis: simulated points of the inverse avidity of CD8 T cells, denoted *I*_*E*0,_ distributed in a way similar to data from patients in another study [2]. Two scaling parameters are used for the two avidities, one for each axis. Red symbols (o) correspond to spontaneous controllers, and blue symbols (o, *) correspond to untreated patients with high viremia and patients on long-term ART, respectively. Blue and red lines show the phase diagrams predicted in this work at two different values of infectivity ratio between activated and resting CD4 T cells, *w* = 100 and *w* = 1. The other parameters are given in Table 1.

Due to complex dynamics of the host-virus system, mathematical modeling has been employed to describe various aspects of HIV infection [35-44], including the existence of progression and control [45] and the treatment with monoclonal antibodies [46]. The relative roles of the fitness cost of antigen escape from CD8 T cell response and the breadth of recognition have also been addressed [47]. However, the striking difference observed in the distribution of the functional avidity of helper CD4 T cells between controllers and progressors, in the absence of such difference for CD8 T cells, remains unexplained [1, 2]. The observation is especially interesting given that genetic markers of CD8 T cell receptors (HLA subtype) remain the strongest predictor of HIV control [21-24].

**Table 1.** Model parameters and their default values in simulation of a representative progressors and controllers. The normal CD4 count, *T*_0_, and the maximal allowed values of *H, E* are set to 1 and used as units for the respective variables: *T, I, L*, and *H, E*. The general ranges of parameters are taken from [43, 44]. The specific (default) values of parameters are adjusted to fit data in Fig. 1 and to obtain oscillation-free dynamics in Figs. 3 and 4, and to match the representative levels of various cell types (see experimental references in [42]).

| Notation | Parameter name | Unit | Default | Range |
| --- | --- | --- | --- | --- |
| $I_{0H}$ | 1/avidity helper cells | $T_0$ | $9 \cdot 10^{-6}$ | $10^{-6} - 1$ |
| $I_{0E}$ | 1/avidity killer cells | $T_0$ | 0.03 | $10^{-6} - 1$ |
| $w$ | H to T infectivity ratio | 1 | 10 | $1 - 100$ |
| $c$ | Max. proliferation rate | 1/day | 2.0 | $1.5 - 3$ |
| $d_I$ | Infected cell death rate | 1/day | 1.0 | $0.7 - 1.2$ |
| $d_T$ | Target cell death rate | 1/day | 0.3 | $0.1 - 0.5$ |
| $d_H$ | Helper cell death rate | 1/day | 0.15 | $0.07 - 0.3$ |
| $d_E$ | Killer cell death rate | 1/day | 0.15 | $0.07 - 0.3$ |
| $r_{max}$ | Max. latency activation rate | 1/day | 0.2 | |
| $r_0$ | Min. latency activation rate | 1/day | 0.001 | |
| $p_{L0}$ | Max. latency probability | 1 | 0.4 | |
| $R_0$ | Basic reproduction number | 1 | 8 | $4 - 12$ |
| $\varepsilon$ | Drug efficacy | 1 | 0.97 | |
| $I_{in}$ | Initial actively infected cell # | $T_0$ | $10^{-11}$ | $10^{-11} - 10^{-9}$ |
| $L_{in}$ | Initial latent cell # | $T_0$ | $10^{-11}$ | $10^{-11} - 10^{-9}$ |
| $E_{in}$ | Initial killer cell # | 1 | $10^{-5}$ | $10^{-7} - 10^{-4}$ |
| $H_{in}$ | Initial helper cell # | 1 | $10^{-5}$ | $10^{-7} - 10^{-4}$ |
| $T_0$ | Normal target cell # | 1 | 1 | |
| $H_0$ | Acting # of helper cells | 1 | $10^{-2}$ | |
| $E_0$ | Acting # of killer cells | 1 | $10^{-2}$ | |

The present work offers a model that explains these findings and the difference in the virus dynamics between controllers and progressors. The starting point is the observation that the avidity of effector CD8 T cells, in terms of the average antigen concentration in the body, is usually much lower than that of effector CD4 T cells, even though both of them span several orders of magnitude. Indeed, effector CD8 T cell proliferates only on a direct contact with an infected cell expressing MHC-I-peptide complex, while an effector CD4 T cell reacts to extra-cellular soluble antigen presented by specialized MHC-II cells and hence can sense antigen far away from its cellular source [48]. Next, HIV and a few other viruses, such as LCMV [49], have the ability to infect and deplete specifically CD4 cells, either by direct cytotoxicity or through killer cells. Taken together, these two observations appear to suggest that the survival strategy of HIV may be to suppress the CD4 T cell response and thus make the immune system much less sensitive to the presence of the virus. Thirdly, it is well known that HIV infects activated CD4 T cells much faster than resting cells [44, 50]. The reason is a much higher concentration of nucleotides in activated cells [51]. Based on these facts, I propose a model, which predicts the infection outcome based on the knowledge of patient’s parameters and explains the difference in CD4 T cells avidity between patient cohorts. The model is compared to data on T-cell avidity from two recent studies [1], [2]. In *Discussion*, we discuss the alternative explanation of the dual outcome of HIV infection based on T cell exhaustion and its connection to experimental data.

## Results

### Model

The model comprises five cell compartments, as follows (Fig. 2): uninfected but susceptible target cells of CD4 T CCR5 phenotype, which number is denoted *T*; actively infected cells, *I*; latently-infected cells, *L*; virus-specific helper CD4 T cells, *H*; and cytotoxic effector CD8 T cells, *E*. The biological processes included in the model are, as follows.

**Fig. 2.**
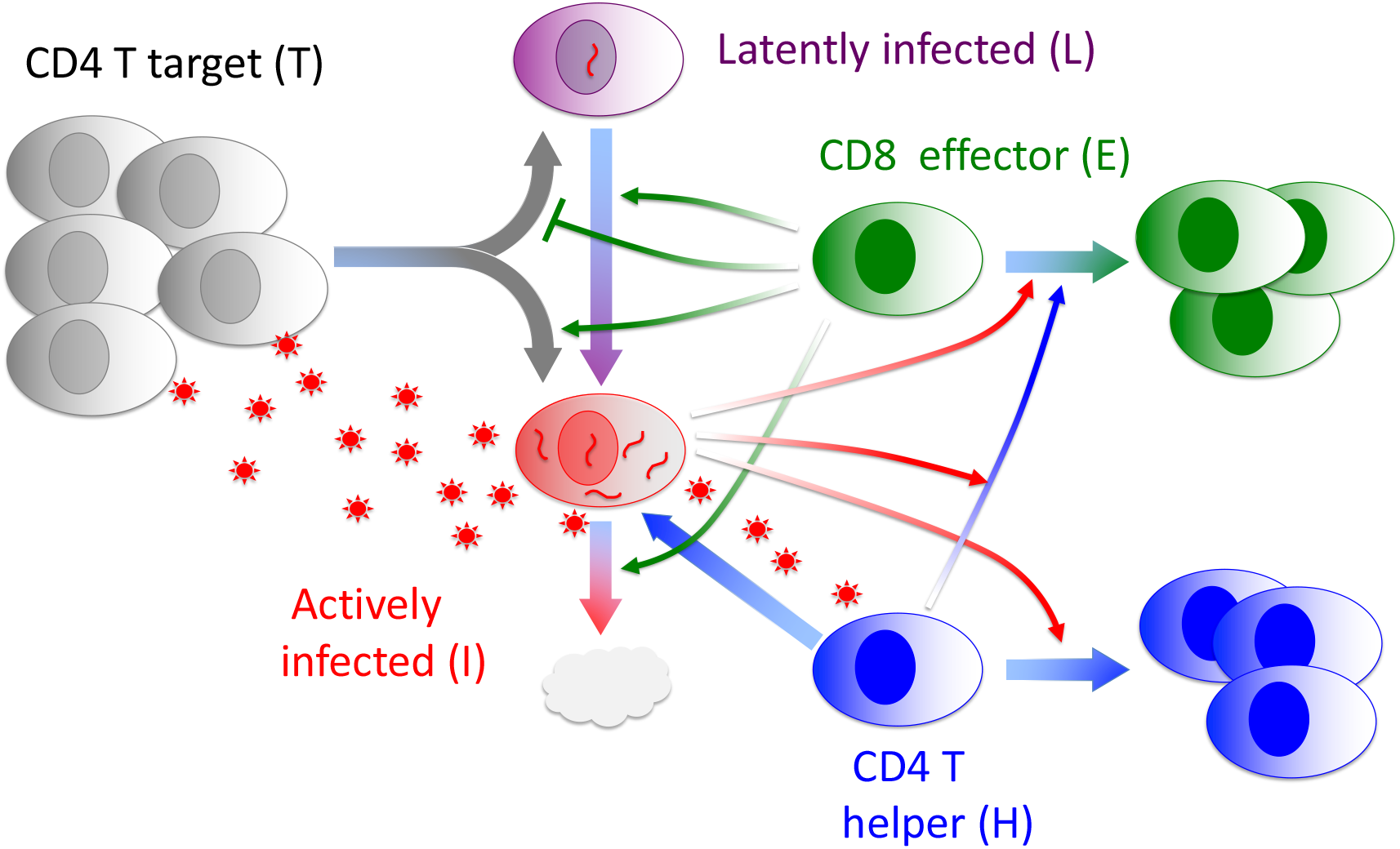
The model of the immune response and virus dynamics explaining the multiple outcomes of HIV infection. Broad arrows show cell flow and proliferation, thin arrows show control signals discussed in the main text. Little red stars are free virus. Model equations are given in *Methods*.

Target CD4 T cells are replenished, are infected with the virus, and have a finite life span. Actively infected cells are expanded due to new infections and reactivation from latently-infected cells and die due to the virus-induced death and killing by cytotoxic effector CD8 T cells. Latently-infected cells are produced upon infection of target cells and can be reactivated, with the rates of both processes controlled by the number of effector CD8 T cells secreting cytokines activating the integrated HIV provirus [52]. HIV-specific CD4 T helper cells proliferate in response to antigen, with a characteristic threshold (inverse avidity) denoted *I*_*H*0_. These cells are infected at a rate higher than non-specific target cells, by a factor denoted *w, w* > 1. The biological justification is that virus-specific cells are activated by antigen, and CD4 T cell infectivity increases with cell activation [44, 50]. CD4 T helper cells also have a finite life span. Finally, HIV-specific CD8 killer cells can proliferate by one of two alternative signaling pathways: either due to binding of TCR to MHC-I-peptide complex expressed by infected cells, with inverse avidity *I*_*E*0_, or due to stimulation of their Il-2 receptor by IL-2 secreted by the helper cells. The model equations are given below in *Methods*. The model parameters including their default values used for simulation are listed in Table 1.

For the reasons detailed in the *Discussion* section, the model does not include the exhaustion of CD8 T cells previously proposed to explain the different outcome in controllers and progressors [45]. In *Discussion* section, we rule out this alternative explanation based on data in Fig. 1 and additional data.

Here I assume that the exhaustion in a chronic infection, which accumulates rather gradually in chronic infection [53], is a consequence of virus persistence rather than its cause, and that its effect on virus dynamics during the first two or three weeks of systemic infection is negligible.

### Two possible steady states

Analysis presented in S1 Text online shows that the system has two possible steady states (fixed points) at which the size of each cell compartment does not change in time because its proliferation is exactly compensated by the cell death, as follows:

1. The *helper-dependent* steady state with a very low virus load observed in a spontaneous or post-treatment controller (*S1 Text*, Eq. 9). An example of simulation resulting in spontaneous control is presented in Fig. 3A, with the stability region is shown in the phase diagram (Fig. 3B, blue region). If the avidity of helper cells is sufficiently low, this state becomes unstable (Fig 3B, green) or disappears altogether (Fig 3B, yellow). The exact conditions of its existence and stability are given in *S1 Text* (Eqs. 15 and 17).
2. The *helper-independent* steady state with a high virus load observed in a progressor or potential post-treatment controller in the absence of treatment (Eq. 10). An example of simulation is shown in Fig. 4A.

**Figure 3.**
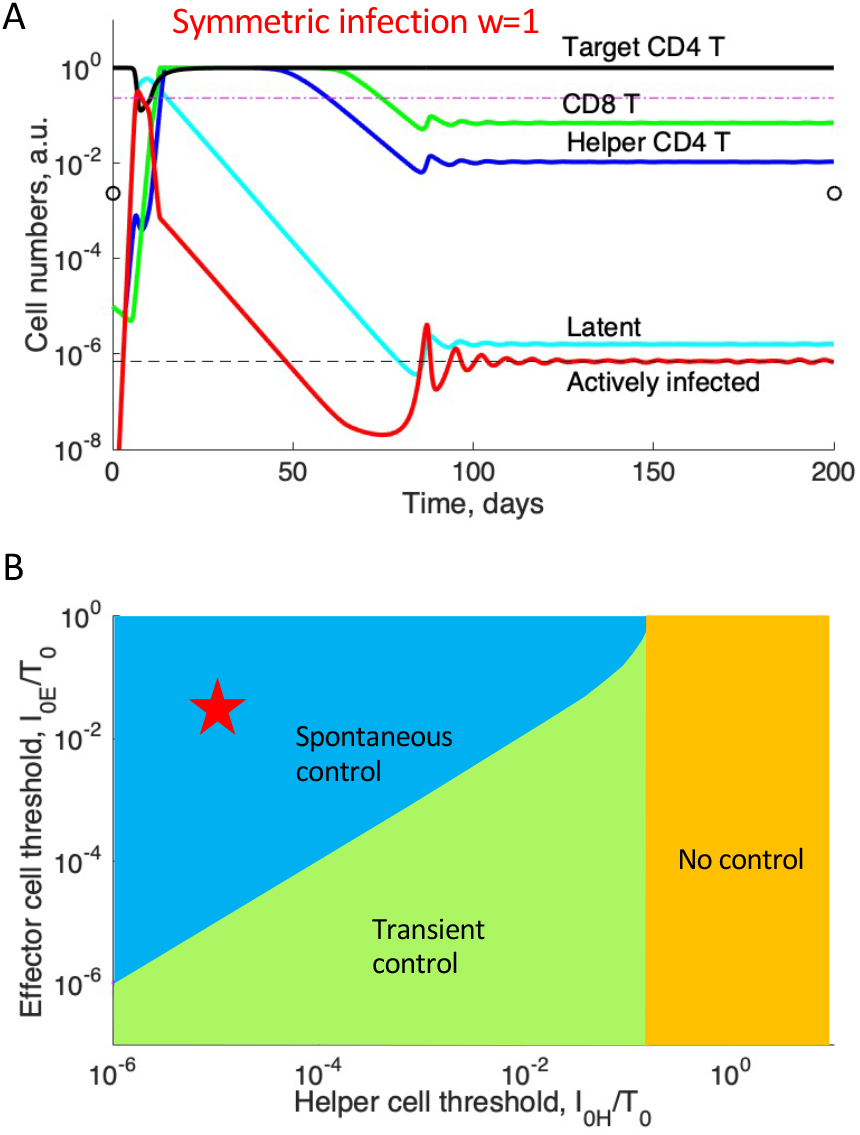
The model predicts the conditions of the spontaneous virus control. Infectivity of virus-specific and non-specific cells is assumed to be the same, *w* = 1. (A) Spontaneous establishment of a low-viremia helper-dependent state. Y-axis: Numbers of cell compartments in relative units to their maximal values set at 1. X-axis: Time of the systemic phase of infection. (B) Phase diagram predicting the four outcomes (S1 Text, Eqs. 15 and 17). X-axis: The helper CD4 T cell threshold in antigen (inverse avidity), *I*_0*H*_, in units of the normal CD4 T cell count, *T*_0_. Y-axis: The same for killer CD8 T cells. The asterisk shows 1/avidities used in (A), *I*_0*E*_ = 0.03, *I*_0*H*_ = 9 ∙ 10^−6^. Notation in (A): Black dashed line is the helper-dependent steady state *I*_1_ (Eq. 9 in S1 Text). Two little circles show the helper-free steady state *I*_2_ (Eq. 10 in S1 Text). Magenta dash-dotted line is the limit of helper cell expansion, *I*^*^, Eq. 11. (A, B) Fixed parameter values are given in Table 1. Relevant parameters in (B): 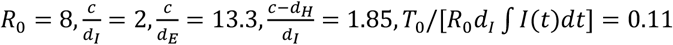. The last parameter is calculated from simulation and depends on the other parameters. The specific (default) values of parameters are adjusted to fit data in Fig. 1 and to obtain oscillation-free dynamics and representative levels of various cell types (see experimental references in [42]).

**Figure 4.**
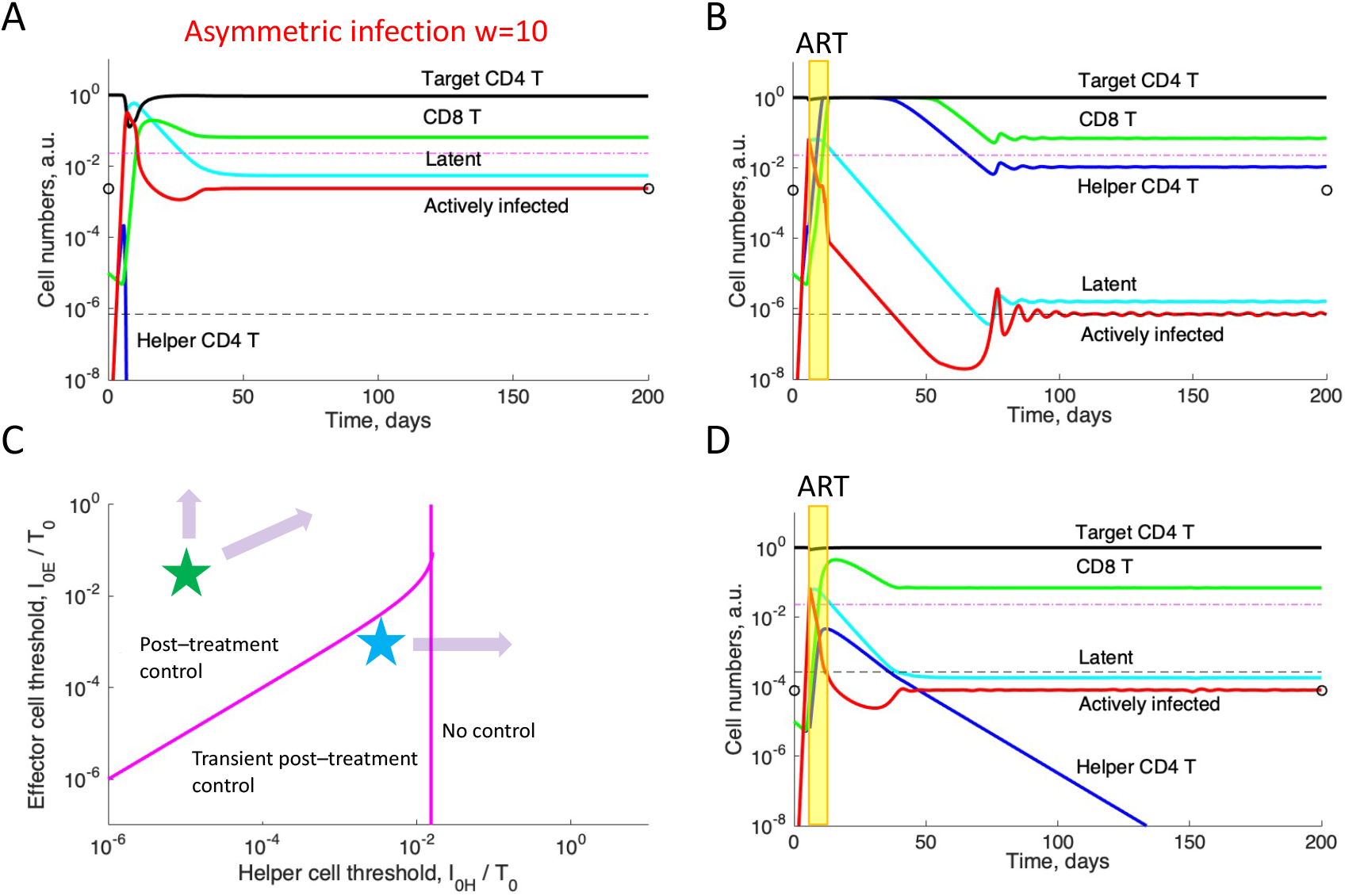
The model predicts the conditions of post-treatment control and transient control. Infectivity of virus-specific (and hence activated) CD4 T cells is assumed to be 10 times higher than for non-specific (and hence resting) CD4 T cells, *w* = 10. (A) Early infection and killing of CD4 T cells leads to a high-viremia helper-free state (*S1 Text*, Eqs. 18, 19). (B) ART between days 6 and 12 with the efficacy of 97% leads to the rescue of helper cells and establishment of a low-viremia state. (D) If avidity of helper cells is lower than the avidity of killer cells, helper cells decay gradually and control is only transient. (A, B, D) Notation as in Fig. 3A. (C) Phase diagram predicting the three outcomes of infection (Eqs. 15, 17). The asterisks show parameters used in simulation: (A,B) *I*_0*E*_ = 0.03, *I*_0*H*_ = 9 ∙ 10^−6^, (D) *I*_0*E*_ = 0.001, *I*_0*H*_ = 0.003. Relevant parameters in (D) are *w* = 10, *T*_0_/ [*R*_0_*d*_*I*_ ∫ *I*(*t*)*dt*] = 0.2 and the other parameters are as in Fig. 3.

### Post-treatment control

Simulation demonstrates that, in the imaginary organism with an unlimited number of T cells, only one of these two states can exist indefinitely, whichever has a lower virus load. In a real organism containing ∼10^11^ T cells, the two states can co-exist. The limited size leads to effective bi-stability, as follows: if the infectivity ratio of active-to-resting helper cells, *w*, is sufficiently high, helper cells can be completely depleted during the acute phase of infection before they can reach the steady state (*S1 Text*, Eq. 19). Even though the helper-dependent state would be stable once reached, helper cells can never make it. As a result, the high-viremia state is established forever (Fig. 4A).

Simulation demonstrates that the helper cell-dependent, low-viremia state can be recovered by an early anti-retroviral therapy (ART) started at day 6 or 7 of systemic infection (that follows the mucosal phase). Blocking infection of helper cells by ART allows them to expand until they reach the steady state and take control of killer cells (Fig. 4B). Because their activation threshold is very low compared to that of killer CD8 cells, the steady state viremia is very low, relatively speaking, which effect explains the effect of post-treatment control.

If the avidity of helper cells is sufficiently small, the post-treatment state will be only transiently stable, helper cells will decay gradually, and the system will eventually slide into the high-viremia state (Fig. 4D).

### Phase diagram of patient cohorts

Combining all these cases (Eqs. 15, 17, and 19 in *S1 Text*, and Fig. 3B and 4C) results in a phase diagram in coordinates (*I*_0*H*_, *I*_0*E*_), with four distinct regions corresponding to four different cohorts, as follows:

- Spontaneous control. A stable helper response and low viremia in the form of oscillations emerge spontaneously (Fig 3A). No treatment is required.
- Post-treatment control. Untreated patients lose helper cells rapidly due to HIV infection (Fig. 4A), but an early antiretroviral treatment rescues helper cells and establishes a low-viremia state (Fig. 4B).
- Transient (unstable) post-treatment control. Helper cells can reach high levels but then decay gradually (Fig. 4D).
- Progressors. The helper-free state with a high virus load is the only possible state, and its dynamics is similar to that in Fig. 4A.

The phase diagram in Fig. 3B and 4C is consistent with the clinical observation that controllers harbor helper CD4 T cells with a higher avidity than progressors [1]. It also demonstrates the importance of an asymmetric infection rate [44, 50].

These results also demonstrate that HIV takes advantage of latency hard-wired into its genetic circuitry [52] to avoid full clearance in controllers by frequent reactivation from latent cells (Fig. 3A and 4B). Another evolutionary role for latency is to improve the chances of viral survival upon mucosal entry [42]. The two evolutionary roles can be combined stating that latency is the method used by HIV to survive population bottlenecks, similar to spores of monocellular organisms [42].

In the phase diagrams (Figs. 3B and 4C), spontaneous controllers differ from post-treatment controllers by an unusually low infectivity asymmetry, *w*. Because *w* is typically quite high [43, 44], post-treatment control (Fig. 4C) is predicted to be much more likely than the spontaneous control (Fig. 3B), which agrees with observations (10-15% and 0.5% of patients, respectively). This prediction is testable, both *in vivo* and *ex vivo*.

An interesting feature is the prediction of a transient controller state, characterized by the slow decay of helper cells and gradual switch to the direct antigen activation (Fig. 3E). This explains, at least, qualitatively, why control is often transient. In principle, control in this state could be prolonged by periodic short-term treatment resetting helper cells to a high level. However, the safety of such a hypothetical protocol, in view of the danger of drug-resistance mutants, would require careful testing.

In addition to predicting the outcome of an individual infection, the model predicts the best timing for ART. Simulation (Fig 4B) demonstrates that ART must start between days 6 and 8 of the systemic (post-mucosal) phase of infection. The exact time window depends on patient’s parameters. The rather strict timing requirement raises the possibility that the observed post-treatment controllers may be only a small fraction of their potential number. In principle, the application of this model could enable physicians to improve the timing of early ART and expand the number of post-treatment controllers. Whether this idea is practical, remains to be seen.

### The model explains the differences in CD4 cell avidity between controllers and progressors

Fig. 1 compares the phase diagram of patient cohorts between two values of infection asymmetry, a high (usual) value, *w* = 100, and a very low value, *w* = 1. Note that the region of virus control is smaller in the first case (blue lines). To test this diagram experimentally, one has to plot on top of it the pairs of avidity values obtained for real patients. Data for the avidity of both T cell types measured in the same study are not available. To partly circumvent this problem, the actual data set on CD4 cell 1/avidity, *I*_*H*0,_ obtained for spontaneous controllers, progressors, and patients on ART in [1] is combined with a simulated set for *I*_*E*0_ resembling experimental ldata for CD8 T cells obtained in [2] (Fig. 1). Based on the result of [2], the same distribution function for the CD8 T cell avidity for all three patient cohorts is assumed. Because both avidities are defined within a constant scaling factor, the two scaling factors are adjusted to fit the data in the diagram; the same two scaling factors are used for all cohorts. Thus, the model explains the observation that progressors have a relatively low CD4 T cell avidity (high *I*_*H*0_).

Spontaneous control is predicted to be a result of two rare events: unusually high CD4 T cell avidity and an unusually low value of *w*. This prediction explains why it spontaneous control so rare (0.5%). In contrast, post-treatment control requires only a high CD4 T cell avidity but asymmetry can by typically high.

### Robustness to parameter values

So far, the focus was on three parameters (*I*_0*H*_, *I*_0*E*_, and *w*) keeping the other model parameters in Table 1 fixed. The sensitivity of the phase diagram to the other parameters is tested with plots made in another pair of axes, (*I*_*H*0_, *w*), while keeping 1/avidity of CD8 T cells, *I*_0*E*_, fixed (*S1 Fig*). The border between progressors and transient controllers is rather robust to all parameters. In contrast, the boundary separating spontaneous control and post-treatment control is sensitive to most parameters. Parameters that most significantly alter the external boundary of post-treatment control are *d*_*H*_ and *d*_*E*_. A particular form of the control functions, *α*(*x*), *β*(*y*) in Eq. 1, is tested not to be very important for results, as long as *α*(*x*) changes between *x* and 1 at small and large *x*, and *β*(*y*) changes between 1 and 0 at small *y* and *y* = 1. The reason for this robustness is that the system has very fast dynamics and spends most of its time at the values of *x* either much larger or much smaller than 1, and at *y* much smaller than 1.

## Discussion

Thus, the model explains why controllers have higher avidity of CD4 T cells than progressors, and explains why post-treatment control is more frequent that spontaneous control. The outcome of HIV infection can be understood from the two signals of CD8 T cell activation, one through T-cell receptor binding to MCH-I-peptide complex of infected cells, and another through IL-2 secretion by virus-specific CD4 T helper cells. The model predicts two possible steady states, one state with a high level of helper cells and a low virus load determined by the maintenance of helper cell population by MHC-II dependent activation, and the other state without helper cells, where the high virus load is self-tuned to maintain the CD8 T cell population by MHC-I dependent activation. The first state is characterized by a very slow progression to AIDS.

Which state is stable, depends, mostly, on three parameters: the avidities of helper cells and CD8 T cells, and the asymmetry of infectivity between antigen-activated and non-virus-specific CD4 T cells. If antigen-specific helper CD4 cells are much more infectable than non-HIV specific CD4 cells (*w* > 2 − 3), which is expected to be the case in most patients, helper cells are completely depleted by virus before they even reach the steady state. As a result, the system arrives at a helper-free state with a high virus load. If, however, a short therapy is administered early, roughly two weeks after infection, infection is transiently suppressed, helper cells are rescued, and the system arrives at the low-viremia state.

Spontaneous control is predicted as a result of two unlikely events: unusually high CD4 T cell avidity and an unusually low value of *w*, in the tail of its distribution. This prediction explains why it is so rare (0.5%). In contrast, post-treatment control requires only a high CD4 T cell avidity but asymmetry can by typically high, which explains why post-treatment control is more frequent that spontaneous control. Thus, not only different outcomes are explained, but also their probabilities and the data separation.

The present model assumed that all CD4 T cells are killed by productive infection. In fact, 95% of T cells die after an abortive infection [54]. There is no need, however to introduce this parameter and repeat simulations again. It suffices to rescale *T*_0_ → 0.05*T*_0_. Indeed, effectively, only 5% of permissive cells become host cells. That means that all the characteristic values of the infected cell number, such as

*I*_0*H*_ and *I*_0*E*_, must be rescaled by the scaling factor of 0.05. Therefore, the indirect death does not change the shape of the phase diagram, as long as *I*_0*H*_ and *I*_0*E*_ are scaled in units of 0.05*T*_0_ (Figs. 2B and 3C). One must use the rescaling factor when comparing them with real data, where *I*_0*H*_ and *I*_0*E*_ are measured in units of cell/ml^8^.

The model predicts a controller state with either flat viremia or periodic oscillations around the steady state value (Fig. 2A or 3B). In fact, viral blips are observed in many patients due to the activation of virus from latent cells modeled earlier in ART patients [55]. This stochastic process is better described by a Gillespie simulation than deterministic equations [42].

The exact timing of early ART is of essence. The best timing can be predicted from the present model informed by the avidities of both T cell types based on the HLA subtypes of the patient. Thus, the present study offers not only a mechanistic method to understand and predict the outcome of HIV infection in an individual, but also a hope to expand the number of post-treatment controllers in the future therapy. Variation of HLA CD8 T cell subtypes remains the strongest genetic determinant of HIV progression [21-24]. Yet, the present model does not predict any interesting dependence on the properties of CD8 T cells. This is because our simple model of immune dynamics (Fig. 2) does not take into account that the virus can mutate in epitopes. In real patients, HIV undergoes up to 20 or 30 primary escape mutations in multiple epitopes, with a widely variable fitness cost [56-60]. The order of these mutations is predicted to be determined by the balance between the partial loss of immune recognition per mutation and its fitness cost [47]. Escape mutations can occur very early in the acute phase, during the peak of infection. This fact limits the longevity of the otherwise stable low–viremia state. For it to be long–lived, an escape from CD8 T cell response controlled by helper cells has to be slow, which requires either a high mutation cost or a small recognition loss per mutation [47]. This requirement may explain why specific HLA subtypes of CD8 T cells correlate with control (see [24] and references therein). In keeping with this prediction, an experimental study from MIT has demonstrated that HLA-B57, the allele the most associated with HIV control, has unusually broad cross-recognition, thus confirming the role of antigenic escape in limiting virus control [25].

Previously, the exhaustion of CD8 T cells has been proposed to explain the difference between controllers and progressors [45]. Citing the evidence of the CD8 T cell anergy in a long-term HIV infection [53, 61], the authors of the model postulate that sufficiently high virus loads can disable virus-specific effector CD8 cells in the acute phase of infection. The acute phase of infection, according to [45], is a battle between the viral expansion that disables CD8 T cells and the expansion of CD8 T cells that kill the infected cells. In patients with a low killing CD8 cell ability, in which case the virus peak is tall enough to suppress the immune response, the virus wins the battle resulting into a high-viremia steady-state. The rare patients with an unusually high killing ability reach high levels of CD8 response with no virus replication in the end, which are the controllers. Finally, patients with an intermediate killing potential are the post-treatment controllers that reach a relatively low viremia after early treatment. The model was also used to interpret the results of treatment of experimentally infected animals with broadly neutralizing antibodies [46] and is now being applied to rare chronic cases of SARS CoV-2. While a valuable attempt at explaining dichotomy “control-progression”, this model has a number of issues regarding its connection to the actual virology and pathogenesis of HIV, as follows:

i. The model [45] postulates instant CD8 T cell exhaustion as a dominant force within the acute infection. In fact, CD8 T cell anergy, which represents a defense mechanism against immunopathology, is associated with long-term chronic infections, including HIV and some LCMV strains [49, 53, 61]. Accumulation of anergic cells in an HIV infection is gradual and progressive [53]. In accordance with this fact, T cell “exhaustion” has been modeled either through a time integral of virus load [62, 63] or through impairment/infection of an additional cell compartment [64].
ii. Given that the outcome (progressive vs control) is decided within the first weeks of infection [11], and that exhaustion is a gradual progressive process [53], it is unclear how the second can be the cause of the first. More likely, it is the other way around.
iii. Further, the cited model makes no attempt to use of virological details specific for HIV of LCMV, which viruses show long-term persistence and exhaustion markers. It does not explain why influenza virus does not persist, despite comparably high peak virus loads. In particular, the model does not take an advantage of the specific tropism of both HIV and LCMV for helper CD4 T cells.
iv. Although CD4 T cell depletion in progressors results in AIDS, the “exhaustion” model fails to recognize CD4 T helper cells as a critically important part of the immune system and does not even include them into consideration [45]. As already mentioned, the avidity of CD4 cells differ drastically between progressors and controllers [1, 65, 66], which fact remains unexplained.
v. The model predicts zero replication in spontaneous controllers [45]. In contrast to the prediction, the active replication levels in spontaneous controllers are never zero, even though some of them are below the detection threshold (1000 to 10,000 actively infected cells in the body) [11].
vi. The exhaustion model predicts that CD8 T cells in controllers have a higher killing capacity than in progressors [45], which, to my knowledge, is not supported by any data. It also predicts a lower viremia peak for controllers than for progressors, which has not been observed either. If anything, controllers have a higher peak. In fact, the controlling HLA subtypes are associated with a broader cross-reactivity [25], which make them less prone to antigenic escape [47].
vii. Although cellular markers of “exhaustion” are known, its molecular mechanisms remain obscure [53], which makes the model difficult to test or falsify.

In conclusion, the present study offers a model of two-signal activation of CD8 T cells that explains the unusually high avidity of helper cells in controllers and predicts a testable connection between HIV control, the avidity of CD4 T cells, and the dependence of HIV infectivity on cell activation status. The last prediction merits further experimental investigation.

## Methods

The time dependence is described by ordinary differential equations for the number of CD4 T cells infectable by virus, *T*, actively infected cells, *I*, latently infected cells, *L*, helper CD4 T cells, *H*, and cytotoxic effector CD8 T cells, *E*, for the model in Fig. 2, have a form

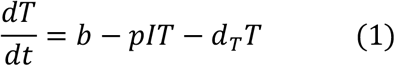

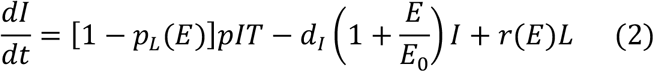

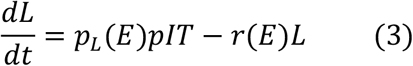

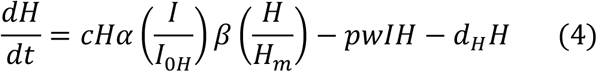

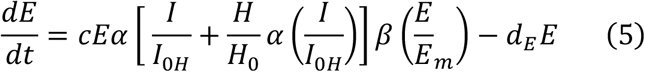

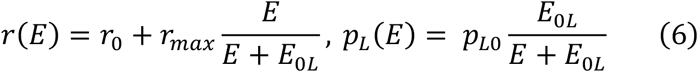

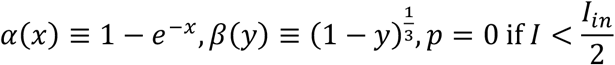

Equation 1 describes the replenishment of target cells (CD4 T+ CCR5+ cells), *T*(*t*), with linear rate *b*, their infection with infectivity parameter *p* and their natural life span 1/*d*_*T*_. Equation 2 describes the expansion and death of actively infected cells, *I*(*t*), as well as their reactivation from latently infected cells, *L*. The death rate includes virus-induced death with rate *d*_;_ and killing by cytotoxic effector CD8 T cells, *E*(*t*). Equation 3 represents the expansion and reactivation of latently infected cells, *L*(*t*), where *p*_*L*_(*E*) is the probability of latency and *r*(*E*) is the probability of reactivation depending on the level of cytokines secreted by effector cells, *E* [42].

Equation 4 states that the dominant subset of CD4 T helper cells recognizing HIV, *H*(*t*), proliferates in response to infected cells with maximal rate *c* but cannot exceed a ceiling, *H*_*m*_. In the absence of activation, helper cells retire at rate *d*_*H*_. The activation-induced death is included in the net proliferation rate, *c*. Helper cells can be infected at rate *wp*, which is larger than the infection rate of target cells, *T*, by a large factor *w*, as given by *w* ≫ 1. The CD4 T cells infectivity can vary by 2-3 orders of magnitude depending on a cell activation status [44, 50]. The concentration of nucleotides is much higher in S phase of the cell cycle than in G1 phase. HIV protein VPR arrests the host cell in G2 phase to optimize nucleotide concentration and thus expedite the reverse transcription of HIV RNA to proviral DNA [51].

Finally, Equation 5 describes the expansion of effector CD8 cells, *E*(*t*), due to two alternative signaling pathways: from infected cells through TCR to MHC-I-peptide binding, and from helper cells through Il-2 stimulation. For the sake of simplicity, I assume that these signals are additive. As I have checked by simulation, the results are not very sensitive to the linear approximation or the form of control functions *α* and *β*. The ranges of model parameters given in Table 1 are based on [44]. Analysis of these equations is given in *S1 Text* online.

### Parameter choice

The aim is to obtain a representative example of dual outcome and fit data in Fig. 1. The general ranges of parameters are taken from [43, 44]. The specific (default) values of parameters are adjusted to fit data in Fig. 1 and to obtain oscillation-free dynamics in Figs. 3 and 4, and to match the representative levels of various cell types (see experimental references in [42]).

## Acknowedgement

The last phase of the study was carried out within the framework of the state assignment of the Federal Agency for Scientific Organizations (FASO Russia: topic no. AAAA-A18-118012290142-9).

## Data and materials availability

The simulation code is available at https://github.com/irouzine/HIVcontrol.

## Supplementary Material

**S1 Fig**. The sensitivity of the phase diagram (*w, I*_0*H*_) obtained from Eqs. 15, 17, and 19 in *S1 Text* to variation of model parameters. The default values are *c* = 3, *I*_0*H*_ = *d*_*H*_ = *d*_*E*_ = 0.5, *I*_0*E*_ = 0.003. The other parameters are as in Table 1, unless shown otherwise.

**S1 Text**. Mathematical appendix.

